# A mouse-specific model to detect genes under selection in tumors

**DOI:** 10.1101/2023.04.12.536653

**Authors:** Hai Chen, Jingmin Shu, Li Liu

## Abstract

Mouse is a widely used model organism in cancer research. However, no computational methods exist to identify cancer driver genes in mice due to a lack of labeled training data. To address this knowledge gap, we adapted the GUST (genes under selection in tumors) model, originally trained on human exomes, to mouse exomes using transfer learning. The resulting tool, called GUST-mouse, can estimate long-term and short-term evolutionary selection in mouse tumors, and distinguish between oncogenes, tumor suppressor genes, and passenger genes using high throughput sequencing data. We applied GUST-mouse to analyze 65 exomes of mouse primary breast cancer models, leading to the discovery of 24 driver genes. The GUST-mouse method is available as an open-source R package on github (https://github.com/liliulab/gust.mouse).

## Introduction

Mouse models are indispensable resources that complement human tissues in cancer research (*1*). In parallel with large-scale sequencing efforts in human cancers, whole exome sequencing and whole genome sequencing of mouse tumors have emerged (*2-5*). Sophisticated algorithms have been developed to identify driver genes in human cancers by integrating mutational patterns, somatic evolution, and other informative features extracted from high throughput sequencing data (*6, 7*). However, no such method is currently available for non-human organisms. Researchers using mouse tumor models often rely on the traditional practice of assuming frequently mutated genes as drivers. But not all recurrent mutations are drivers; and hotspot mutations in passenger genes have been reported (*8-10*). Since mouse tumor models are often induced or genetically engineered from specific mouse strains, the high mutation rate and low genetic diversity inevitably result in many shared passenger mutations (*4*). Advanced tools are needed to go beyond mutation frequency to identify *bona fide* drivers in mice.

Supervised machine learning has been widely used to build models for cancer driver gene prediction (*11*). However, unlike human data, which has carefully curated driver genes and passenger genes available for training a supervised model (*12*), the lack of labeled benchmark genes in mice makes it impractical to train a *de novo* classifier. To address this challenge, transductive transfer learning, a technique that adapts a classifier trained on labeled data in the source domain to unlabeled data in the target domain, may be employed (*13*). Transfer learning is suited for knowledge transfer when the source domain and target domain are similar (*14*). Given that the fundamental mechanisms of neoplastic development are conserved in human and mice (*15*), predictive models built on human genes may be leveraged to develop models for mouse.

We have previously developed the GUST (genes under selection in tumors) method that distinguishes oncogenes (OGs), tumor suppressor genes (TSGs), and passenger genes (PGs) in human cancer genomes (*7*). GUST has two functionalities. Firstly, it estimates key parameters that characterize long-term species evolution and short-term tumor evolution, which can be used to prioritize biomarkers (*16, 17*). Secondly, it includes a random forest classifier that predicts cancer driver genes based on the evolutionary parameters and mutation distribution features. Transductive transfer learning to adapt random forest models can be achieved through structure reduction, which progressively prunes the trees (*18*), and through threshold shifting, which adjusts the cutoff value used at each split (*19*). In this study, we present the GUST-mouse method that is adapted from the GUST method using these two algorithms. Application of this new method to mouse exomes of induced breast cancer models revealed known and novel cancer drivers.

### The GUST-mouse Method

#### Source and target domains

The random forest model in the GUST method is trained to classify human genes into OGs, TSGs, and PGs. The source domain for this model consists of 533 labeled human genes (hBenchmark), which were obtained from the published supplementary materials. The target domain is the mouse exome data from a published study of mouse models of breast cancer (*20*). This dataset includes 65 mouse exomes of primary breast cancers (mmBRCA). We downloaded the VCF files from the NCBI GEO database (GSE142387) and extracted somatic mutations in each sample. The human and mouse reference genomes used in this study were GRCh38 (hg38) and GRCm38 (mm10), respectively.

#### Estimating parameters of long-term species evolution for mouse genes

The GUST method utilized the Multiz alignments of protein sequences from 100 vertebrates to compute position-specific evolutionary rates (*21*). Since the Multiz alignments use human as the reference species, we swapped the mouse sequence with the human sequence, removed sites where the mouse sequence contained a gap, and verified that the resulting mouse sequences were identical to those in the mm10 genome. The evolutionary rate (r) at each position was then computed using the Fitch method (*22*), expressed as the number of substitutions per billion years (s/bys).

#### Estimating parameters of short-term somatic evolution in mouse tumors

For each protein-coding gene in the mm10 genome, GUST-mouse simulated saturated point mutations to infer the expected mutational patterns, considering factors such as codon usage, mutation types, and varying mutational rates. Synonymous mutations were used as the neutral baseline. When analyzing a gene that is mutated in a set of mouse tumors, GUST-mouse compares the observed mutation patterns with the expected patterns to infer selection coefficients of missense mutations (ω) and protein-truncating mutations (φ). The inference is obtained using the maximum likelihood estimation procedure implemented in the GUST program (*7*).

#### Extracting features describing mutation distribution

The GUST program captures the mutational profile of a gene using several features including fractions of missense mutations and protein-truncating mutations, size of clusters of mutations forming hotspots, and length of truncated peptides. The GUST program’s functions for calculating these features can be used to analyze and characterize mutational patterns in mouse genes from exome sequencing data..

#### Refining the random forest classifier

The random forest model in GUST uses 10 predictors, including two long-term evolutionary parameters, two short-term evolutionary parameters, and six mutational distribution parameters, to classify genes into OGs, TSGs, and PGs. For a given mouse gene with mutations observed in a set of cancer exomes, GUST-mouse calculates the values of these predictors. Although the class labels of mouse genes are unknown, it is reasonable to assume that genes with similar roles in tumorigenesis tend to cluster together based on the values of these predictors. GUST-mouse then calculates pairwise Euclidean distances (*D*) between mouse genes based on the predictor values and examines how these distances change in each node of the tree. This allows GUST-mouse to refine the classifier based on the patterns of similarity or dissimilarity between genes in the tree nodes, even in the absence of known class labels for the mouse genes.

Given a bifurcating decision tree *T*_*h*_ in the random forest classifier *RF* trained on human data, GUST-mouse implements two types of transductive transfer learning. The first type involves pruning the tree via structure reduction (*18*). Specifically, it traverses the *T*_*h*_ tree from root to leaves in a depth-first order. At each internal node, GUST-mouse calculates the mean distance between all pairs of genes reaching that node (*D*_*i*_), as well as between all pairs of genes reaching each of its child nodes (*D*_*a*_ and *D*_*b*_). If splitting the internal node into the child nodes does not reduce the pairwise gene distance (i.e., *D*_*i*_ < *D*_*a*_ and *D*_*i*_ < *D*_*b*_), the clade below the internal node is snipped. This process is applied recursively to the entire tree, resulting in an updated tree *T*_*prune*_.

The second type of transductive transfer learning in GUST-mouse does not change the topology of the tree, but rather adjusts the splitting threshold of each internal node (*19*). Similar to the first type, GUST-mouse traverses the *T*_*h*_ tree from root to leaves in a depth-first order. At each internal node, the optimal threshold of the splitting feature is selected to minimize the sum of pairwise gene distance in the two child nodes (i.e., *argmin*_*t*_*(D*_*i*_ *+ D*_*a*_) where *t* is the splitting threshold). After completing the traversal and threshold adjustment, the updated tree *T*_*shift*_ is obtained.

The GUST model consists of 200 *T*_*h*_ trees. By applying structure reduction and threshold adjustment to each tree, the GUST-mouse model will have 200 *T*_*prune*_ trees and 200 *T*_*shift*_ trees. These updated trees collectively constitutes the random forest classifier *RF-mouse*.

## Results

### Human genes and mouse genes showed similar distributions of evolutionary parameters

For each gene in the hBenchmark dataset and in the mmBRCA data sets, we calculated the evolutionary rates of each affected position, recurrently mutated positions, and all positions. Low evolutionary rates indicate strong purifying selection across species and large functional impact. The comparison of evolutionary rates between the mmBRCA data set and the hBenchmark data sets (human) showed highly similar distributions (**Fig. 1A**). This suggests that the evolutionary constraints and functional impact of mutations in these datasets are comparable between human and mouse, despite the species differences.

**Figure 1.**
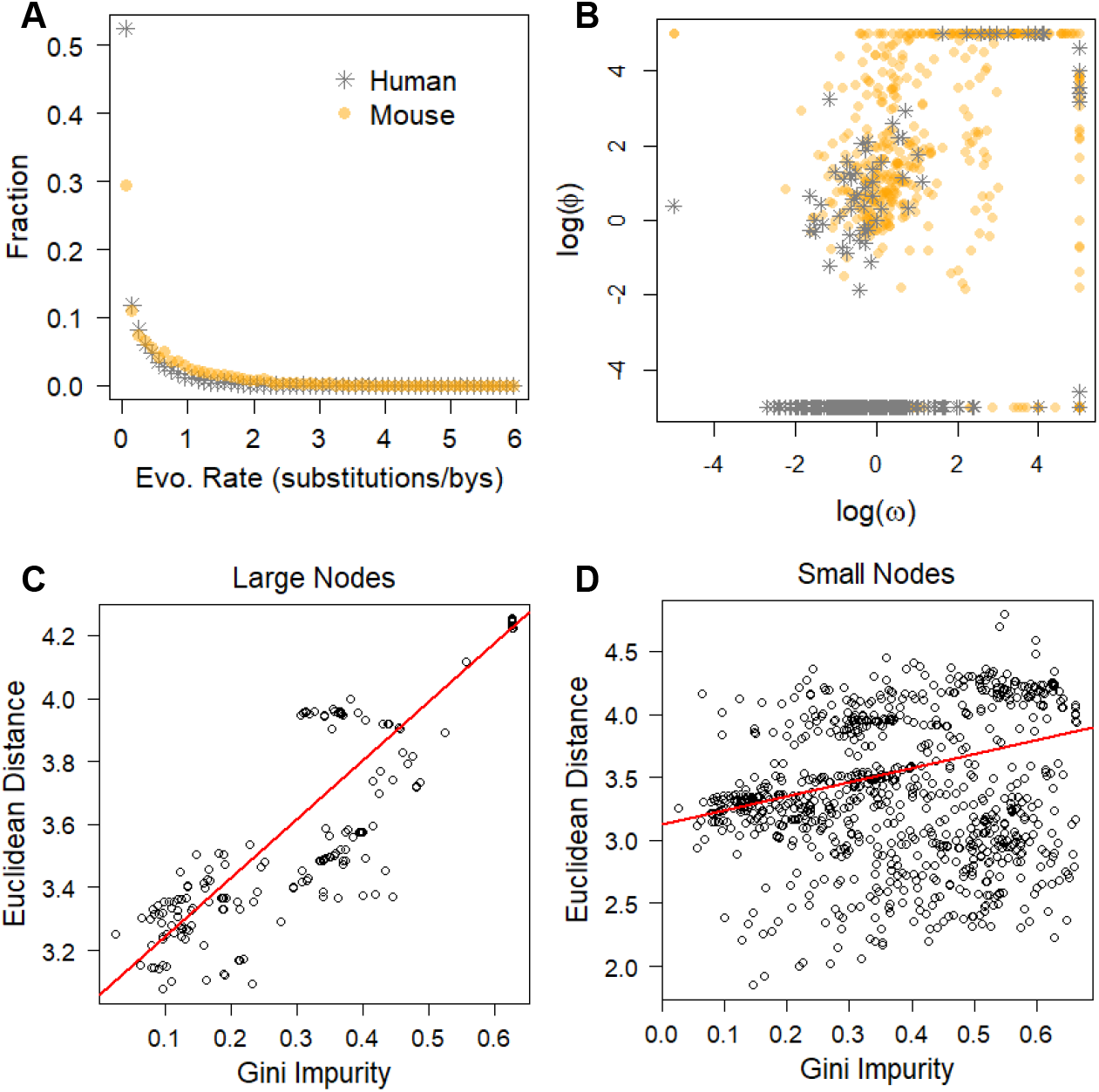
Building the RF-mouse classifier via transfer learning. (**A**). Mutations in the hBenchmark dataset and in the mmBRCA data set showed similar distributions of long-term evolutionary rates. (**B**). Scatterplots of short-term somatic selection of missense mutations measured by log(ω) and truncating mutations measured by log(φ), showed similar distributions in the hBenchmark dataset and in the mmBRCA dataset. (**C, D**) Gini impurity score and within-node Euclidean distance were strongly correlated in large nodes with size > 200 (C) and were moderately correlated in small nodes wit size <20 (D). Red lines represent linear fits.

The selection coefficients, ω and φ, quantify short-term somatic selection on missense mutations and protein-truncating mutations, respectively. The sign of the coefficient indicates direction of selection (positive or negative) and the magnitude indicates strength of selection. We computed ω and φ for mutated genes in the mmBRCA dataset and the hBenchmark dataset. The scatterplots showing the distribution of ω and φ in each dataset shared a similar pattern – most genes were under neutral selection (ω and φ close to 0 on log scale), and a small number of genes were under directional selection (ω and φ deviated from 0, **Fig. 1B**).

The consistent patterns of long-term and short-term evolutionary parameters across data sets confirmed that human and mouse cancers share common molecular mechanisms. This supports the notion that findings from one species can be informative and relevant for understanding cancer biology in the other species and justifies the use of transfer learning approaches.

### Unsupervised Euclidean distance was a good proxy of supervised splitting index

The *RF* model contained 200 *T*_*h*_ trees trained on the labeled hBenchmark data representing the source domain. We previously reported that this model had a cross-validation accuracy of 92% and area under the receiver operating characteristic curve (AUROC) of 0.97 (*7*). In the training of the *RF* model, Gini impurity score was used as the splitting index. To assess if within-node Euclidean distance calculated without knowing class labels was a good proxy of Gini impurity score, we examined each split where a parent node was divided into two child nodes. In 99.6% (1,800 out of 1,807) of the splits, the mean pairwise distance of genes in a child node was smaller than that in the parent node. This observation is consistent with the expectation that node splitting creates clusters of similar genes. Furthermore, the within-node distance was positively correlated with the Gini impurity; and the correlation was stronger in large nodes close to the root than in small nodes close to the leaves (Pearson correlation coefficient range from 0.88 to 0.31, all P<10^−16^, **Fig. 1C-D**). This result confirmed our assumption that genes with different class labels form clusters that can be inferred from Euclidean distance.

### Random forest classifier adapted via transfer learning predicted known and novel driver genes

The mmBRCA data set contained a total of 18,454 somatic mutations (point mutations and short indels) in 1,004 genes. Using these data as the target domain, we adapted the *RF* model and built the *RF-mouse* model. The new model predicted 23 OGs and 1 TSGs with high confidence (probability > 0.95, **Table 1**). The *Cdk12* gene, a known OG and a potential drug target for breast cancer is a representative example (*23*). As expected, missense mutations in this gene were clustered in a hotspot and under strong positive selection (log(ω)=5.0, **Fig. 2A**). In the *Foxn2* gene, a known TSG (*24*), nonsense and frameshifting mutations that truncated the protein and removed the DNA-binding domain were under strong positive selection (log(φ)=3.7, **Fig. 2B**). Therefore, these two genes were putative drivers that confer a selective advantage to cancer cells and promote oncogenesis. Meanwhile, an overwhelming majority of the mutations were predicted as passengers where protein-changing mutations were under similar neutral selection as synonymous mutations, such as the *Gm8909* gene (log(ω)=0.19, log(φ)=0.60, **Fig. 2C**). These findings highlight the ability of the *RF-mouse* model to predict drivers in mouse tumors and provide insights into the functional impact and selection pressures acting on somatic mutations in specific genes.

**Table 1.**
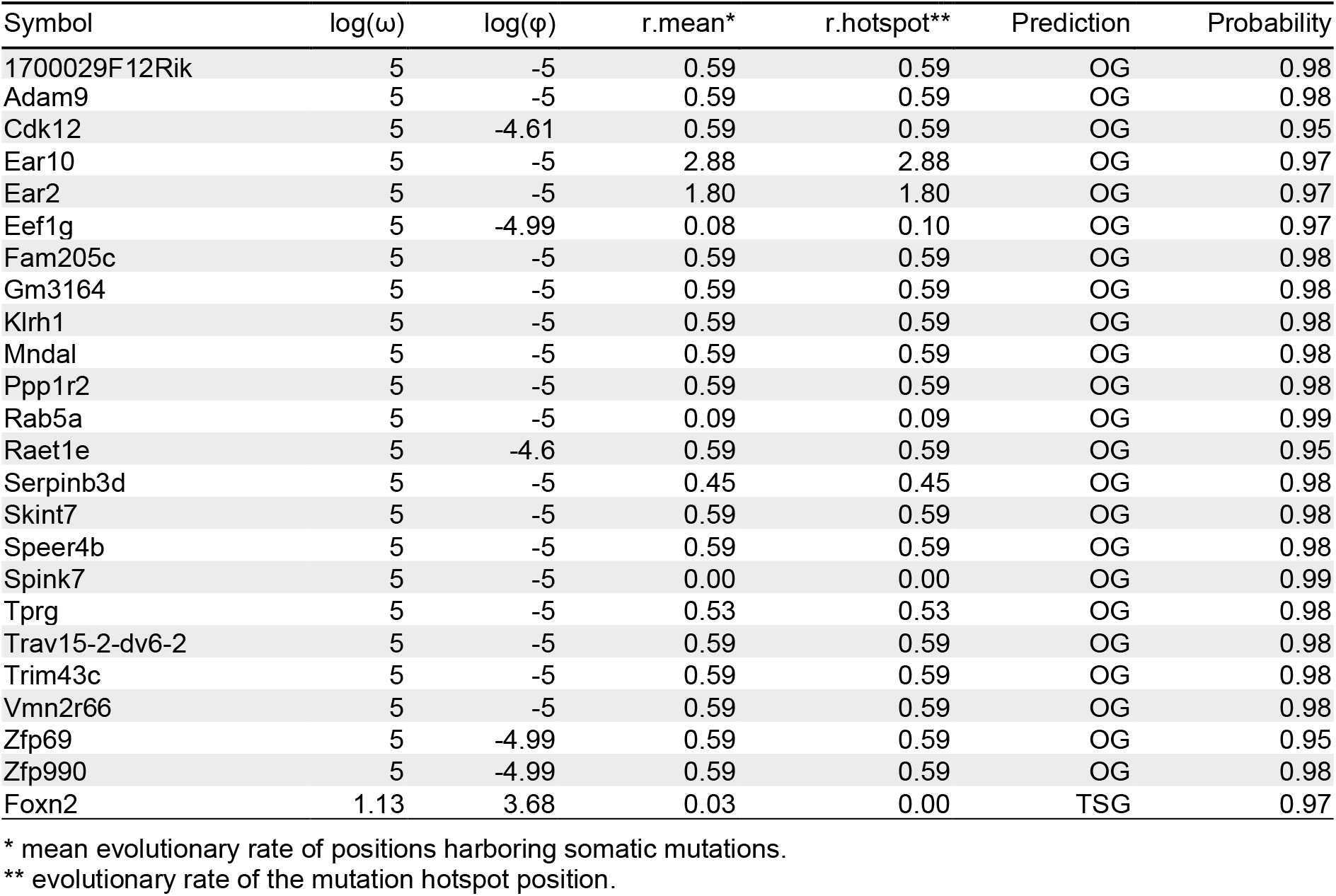
OGs and TSGs predicted with high confidence (probability >0.95)

**Figure 2.**
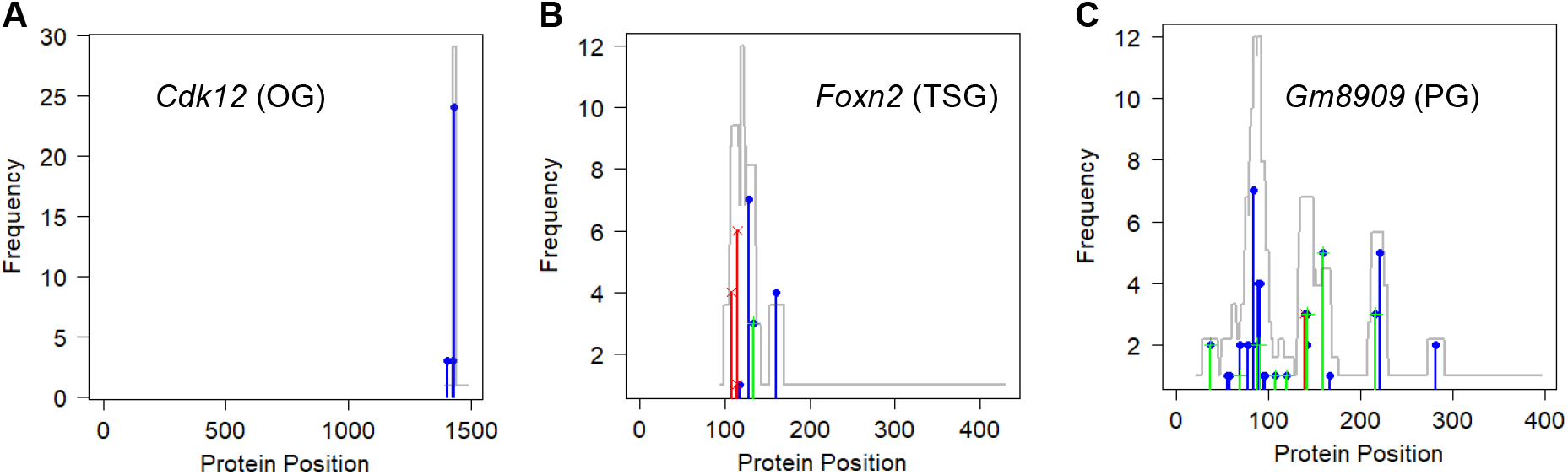
Mutation profiles of representative genes. (**A**) The *Cdk12* gene was predicted as an OG, showing a hotspot of missense mutations (blue bars). (**B**) The *Foxn12* gene was predicted as a TSG, showing a cluster of protein-truncating mutations (red bars) removing peptide after position 115. (C) The *Gm8909* gene was predicted as a PG, with missense (blue bars) and synonymous mutations (green bars) scattered throughout the protein. Mutation density was displayed as gray lines.

## Discussion

Tumorigenesis is an evolutionary process, in which selectively advantageous mutations accumulate in cancer cells, leading to uncontrolled cell growth and tumor formation (*25, 26*). The newly developed computational tool, GUST-mouse, is the first of its kind to enable the study of mouse tumors within an evolutionary framework. It provides two levels of analysis – estimation of evolutionary parameters and classification of driver genes.

From the long-term evolutionary perspective, cancer driver mutations are under strong purifying selection across species and tend to affect highly conserved sites in the genome (*27*). GUST-mouse, through its ability to compute substitution rates at positions affected by different types of somatic mutations, provides valuable information about the evolutionary conservation of mutated sites. Similarly, from the short-term evolutionary perspective, mutations that result in gain-of-function or loss-of-function effects are under strong positive selection, as measured by the selection coefficients (ω and φ). These quantitative measures can assist researchers in biomarker selection and understanding the molecular mechanisms underlying cancer development. Other computational methods can also incorporate these values as prior knowledge into algorithm design.

Due to the scarcity of curated cancer drivers in mice, GUST-mouse relies on transfer learning to adapt the classifier trained on labeled human data to fit in the mouse domain. An important consideration in transfer learning is the similarity between the source domain and the target domain. Theoretically, tumorigenesis in human and in mice share common hallmarks (*15*). Our empirical analysis confirmed that the human and mouse exome data indeed shared similar distributions (**Fig. 1**). Using this adapted classifier, we identified known cancer drivers and passengers with patterns consistent with expectations (**Fig. 2**).

However, models constructed from unlabeled data may have intrinsic weaknesses. While the current literatures report structure reduction and threshold shifting are effective techniques to transfer a random forest model, we were unable to evaluate the performance of the GUST-mouse classifier. To provide transparency to users, GUST-mouse displays a warning message of “accuracy unknown” in the header of the prediction result file. This serves as an alert to users to interpret the results with caution, considering the potential uncertainties. Further research and validation using labeled data in the target domain may be necessary to assess and improve the performance of the GUST-mouse classifier.

We implemented the GUST-mouse as a R package that is freely available on github (https://github.com/liliulab/gust.mouse). Detailed documentation is provided in the standard R manual format.

## Conclusions

The GUST-mouse method provides a mouse-specific model to study long-term and short-term evolution of cancer mutations, and to identify driver genes. It is a valuable computational tool that can contribute to our understanding of tumorigenesis and facilitate comparative studies between human and mouse tumors.

## References

1. K. K. Frese, D. A. Tuveson, Maximizing mouse cancer models. Nat Rev Cancer 7, 645–658 (2007).

2. S. Sneddon et al., Whole exome sequencing of an asbestos-induced wild-type murine model of malignant mesothelioma. BMC Cancer 17, 396 (2017).

3. L. Wei et al., Exome sequencing analysis of murine medulloblastoma models identifies WDR11 as a potential tumor suppressor in Group 3 tumors. Oncotarget 8, 64685–64697 (2017).

4. M. Dow et al., Integrative genomic analysis of mouse and human hepatocellular carcinoma. Proc Natl Acad Sci U S A 115, E9879–E9888 (2018).

5. C. L. Lee et al., Whole-Exome Sequencing of Radiation-Induced Thymic Lymphoma in Mouse Models Identifies Notch1 Activation as a Driver of p53 Wild-Type Lymphoma. Cancer Res 81, 3777–3790 (2021).

6. J. Lyu et al., DORGE: Discovery of Oncogenes and tumoR suppressor genes using Genetic and Epigenetic features. Sci Adv 6, (2020).

7. P. Chandrashekar et al., Somatic selection distinguishes oncogenes and tumor suppressor genes. Bioinformatics 36, 1712–1717 (2020).

8. R. Buisson et al., Passenger hotspot mutations in cancer driven by APOBEC3A and mesoscale genomic features. Science 364, (2019).

9. J. M. Hess et al., Passenger Hotspot Mutations in Cancer. Cancer Cell 36, 288–301 e214 (2019).

10. V. Trevino, HotSpotAnnotations-a database for hotspot mutations and annotations in cancer. Database (Oxford) 2020, (2020).

11. S. Khalighi, S. Singh, V. Varadan, Untangling a complex web: Computational analyses of tumor molecular profiles to decode driver mechanisms. J Genet Genomics 47, 595–609 (2020).

12. P. A. Futreal et al., A census of human cancer genes. Nat Rev Cancer 4, 177–183 (2004).

13. S. J. Pan, Q. Yang, A survey on transfer learning. IEEE Transactions on knowledge and data engineering 22, 1345–1359 (2010).

14. A. Arnold, R. Nallapati, W. W. Cohen, in Seventh IEEE international conference on data mining workshops (ICDMW 2007). (IEEE, 2007), pp.77–82.

15. A. T. Kho et al., Conserved mechanisms across development and tumorigenesis revealed by a mouse development perspective of human cancers. Genes Dev 18, 629–640 (2004).

16. X. Guan, G. Runger, L. Liu, Dynamic incorporation of prior knowledge from multiple domains in biomarker discovery. BMC Bioinformatics 21, 77 (2020).

17. J. Garcia-Pelaez, R. Barbosa-Matos, I. Gullo, F. Carneiro, C. Oliveira, Histological and mutational profile of diffuse gastric cancer: current knowledge and future challenges. Mol Oncol 15, 2841–2867 (2021).

18. N. Segev, M. Harel, S. Mannor, K. Crammer, R. El-Yaniv, Learn on source, refine on target: A model transfer learning framework with random forests. IEEE transactions on pattern analysis and machine intelligence 39, 1811–1824 (2016).

19. X. Liu et al., in Proceedings of the IEEE conference on computer vision and pattern recognition. (2013),pp. 492–499.

20. C. Ross et al., The genomic landscape of metastasis in treatment-naive breast cancer models. PLoS Genet 16, e1008743 (2020).

21. R. M. Kuhn, D. Haussler, W. J. Kent, The UCSC genome browser and associated tools. Brief Bioinform 14, 144–161 (2013).

22. L. Liu, S. Kumar, Evolutionary balancing is critical for correctly forecasting disease-associated amino acid variants. Mol Biol Evol 30, 1252–1257 (2013).

23. H. Liu, K. Liu, Z. Dong, Targeting CDK12 for Cancer Therapy: Function, Mechanism, and Drug Discovery. Cancer Res 81, 18–26 (2021).

24. H. Ye, M. Duan, FOXN2 is downregulated in breast cancer and regulates migration, invasion, and epithelial-mesenchymal transition through regulation of SLUG. Cancer Manag Res 11, 525–535 (2019).

25. N. Ahmadinejad et al., Accurate Identification of Subclones in Tumor Genomes. Mol Biol Evol 39, (2022).

26. M. Casas-Selves, J. Degregori, How cancer shapes evolution, and how evolution shapes cancer. Evolution (N Y) 4, 624–634 (2011).

27. S. Kumar, J. T. Dudley, A. Filipski, L. Liu, Phylomedicine: an evolutionary telescope to explore and diagnose the universe of disease mutations. Trends Genet 27, 377–386 (2011).

